# Cell-scale confinement reorders bacterial speed rankings

**DOI:** 10.64898/2026.11.04.764272

**Authors:** Yoshiki Y. Shimada, Naoki A. Uemura, Aoba Yoshioka, Daisuke Nakane, Tetsuo Kan

**Author notes:** Corresponding author (DN), (TK).

## Abstract

Which bacterium swims fastest depends on where the race is run. Speeds are usually measured in open liquid, yet many bacteria live in pores, mucus and host passages no wider than a cell. Using a microfluidic device with eight channel widths (0.80–3.00 µm) in one field of view, we tracked unlabeled cells of eight species spanning five motility systems. Chamber height, a polymer additive and channel width each reordered the ranking. *Vibrio alginolyticus*, the fastest in open liquid, slowed to ∼4 µm/s in 1.25-µm channels, whereas *Campylobacter jejuni* and *Bacillus subtilis* kept 33–46 µm/s. Open liquid speed did not predict robustness to confinement. In two unrelated lineages, host symbionts that pass through narrow passages during colonization, *Caballeronia insecticola* and *Vibrio fischeri*, outran their non-symbiotic relatives in the channels, and crossed them with flagella wrapped around the cell body at nearly the speed of unwrapped cells. Where space is tight, the ability to keep moving, not top speed, sets the ranking.

## Introduction

Bacteria are usually compared by how fast they swim in open liquid^1,2^. Comparing microbes as competitors has become a popular framing, from the World Cell Race for eukaryotic cells to the Microbial Olympics^3–5^. But a race result depends on the track. The habitats in which bacteria actually move, such as soil pores, mucus, biofilms and host tissues, contain spaces as small as a single cell^6^. Whether the speed ranking of bacterial species measured in open liquid survives cell-scale confinement is therefore an open question.

At the scale of a bacterium, swimming is governed by viscosity rather than inertia^7^. Propulsion comes from the flagellar motor, a rotary machine whose output depends on load^8^, and motors differ among species^9^. Near a wall, drag increases and trajectories bend, so that *Escherichia coli* swims in circles on a surface and keeps to one side of a channel^10^. *E. coli* remains motile in channels about as wide as the cell, where contact with the walls rather than fluid drag appears to limit movement^11^, and in channels a few times wider than the cell it switches to a stable one-dimensional mode of swimming that is at least as fast as swimming along a flat surface^12^. Theory predicts that such confinement can either enhance or hinder propulsion depending on the geometry and gait of the swimmer^13^. Polymer solutions can raise or lower swimming speed depending on the polymer and on the arrangement of the flagella^14,15^. Because bacteria use very different motility systems, including a single polar flagellum, a polar bundle, peritrichous flagella, bipolar flagella and gliding without flagella^16,17^, each species may respond differently to the same physical constraint.

Most of this knowledge comes from one species at a time, usually *E. coli.* Microfluidic and nanofabricated devices now expose bacteria to geometries that approach the size of a cell^18–20^, and single-species studies have shown wall-guided trajectories, trapping in porous media, and changes in swimming and escape behavior in confined spaces^12,21–23^. Only a few studies have compared multiple species. One examined five flagellated species in channels 2–8 µm wide, mostly wider than the cells, and found that the pattern of movement depended on cell shape and channel geometry, whereas speed changed little with width for most species^24^. We recently found that, in 1-µm passages, some polarly flagellated species keep moving whereas others nearly stall^25^. What is still missing is a direct comparison of many motility systems, including gliding, in channels that approach the cell diameter with all widths recorded at once and compared with open liquid. Only such a comparison can test whether the ranking itself is stable.

The question takes a specific form for bacteria that must pass through narrow passages inside their hosts. The insect symbiont *Caballeronia insecticola* passes through a constricted region of the bean bug midgut to reach its symbiotic organ^26–28^, and the squid symbiont *Vibrio fischeri* (*Aliivibrio fischeri*) passes through micrometer-scale ducts and a bottleneck to reach the light organ of its host^29,30^ (Fig. 1a). Both species can swim with their flagella wrapped around the cell body^31,32^, a mode called flagellar wrapping^33^ (Fig. 1b). Wrapping has been found in polarly flagellated bacteria from several lineages, including *Shewanella*, *Pseudomonas*, *Helicobacter, Vibrio*, and *Burkholderia* sensu lato, particularly in viscous or structured environments^31,32,34–37^, and hydrodynamic simulations predict that a wrapped filament propels the cell less efficiently than an extended one in open fluid but more efficiently in narrow tubes^38^. The transition to the wrapped state has been explained as a buckling instability driven by motor torque^39^, and we recently showed that wrapping gives *C. insecticola* a mechanical advantage in a one-micrometer passage and contributes to successful colonization^25^. Whether narrow passages favor symbionts over their non-symbiotic relatives has not been tested. Each symbiont has a close relative that does not form the corresponding symbiosis^40,41^, which allows a direct test. If the narrow passages of hosts have acted as a filter on motility, the symbionts should move better under confinement than their relatives, whatever the mechanism.

**Figure 1:**
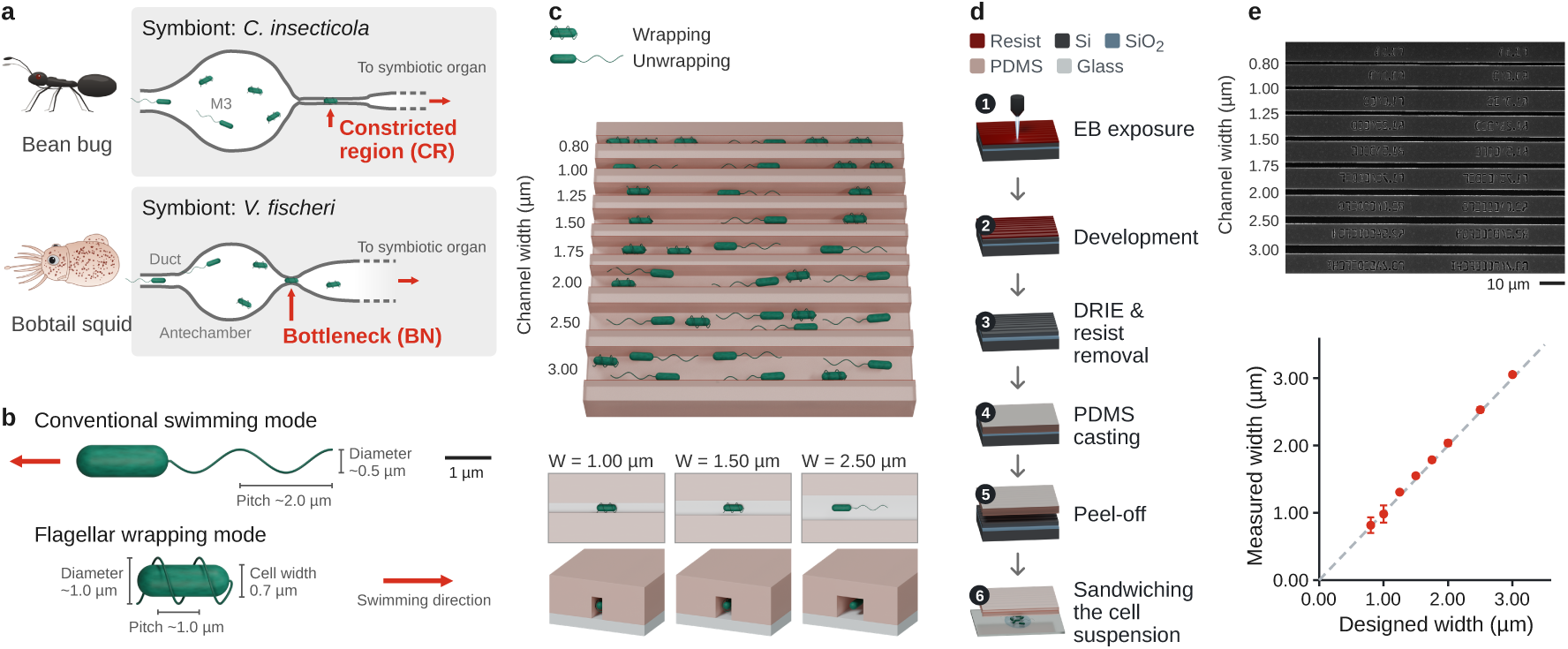
Design and fabrication of a microfluidic platform for bacterial motility phenotyping under cell-scale confinement. **a** Examples of cell-scale confinement encountered by bacterial symbionts in host environments. Schematics illustrate the constricted region of the bean bug gut and the bottleneck of the squid light organ, through which their respective symbionts, *C. insecticola* and *V. fischeri*, migrate. **b** Schematic comparison of conventional swimming and flagellar-wrapping motility. In conventional swimming, flagellar filaments extend behind the cell body, whereas in wrapped-mode swimming, the filaments wrap around the cell body. Representative dimensions of the cell body and flagellar helices are indicated. **c** Design of the microbial confinement assay. Eight parallel microchannel widths, ranging from 0.80 to 3.00 µm, were integrated into a single device, enabling simultaneous observation of bacterial motility under different degrees of confinement within a single microscopic field of view. The channel height was fixed at 2 µm. Enlarged schematics illustrate the channel geometries in top and cross-sectional views. **d** Fabrication process of the microfluidic device. Microchannel patterns were defined on a silicon-on-insulator wafer using electron-beam lithography and dry etching. The resulting silicon mold was replicated in PDMS, and the PDMS device was assembled against a glass surface to confine bacterial cells within the microchannels (see more details in Materials and Methods). **e** SEM image of the fabricated microchannels (upper) and comparison between designed and experimentally measured channel widths (lower; mean ± SD, *N* = 9 channels).

Here we addressed both questions with one device. Parallel channels of eight widths (0.80–3.00 µm; height 2 µm) were placed side by side, all within a single field of view, and an image-processing step recovered cell trajectories without fluorescent labels. Using eight bacterial species with five types of motility system, we compared their performance in open chambers of different heights, with and without a polymer additive, and in channels as narrow as the cell diameter. The ranking changed with every change of environment. The fastest species in open liquid lost more than 90% of its speed under confinement, both host symbionts outran their non-symbiotic relatives in narrow channels, and direct imaging of flagella showed that both symbionts passed through the channels with their flagella wrapped around the cell body, at speeds close to those of unwrapped cells.

## Results

### A microfluidic device for comparing motility under cell-scale confinement

To compare bacterial motility under confinement, we fabricated parallel channels with a fixed height of 2 µm and eight widths from 0.80 to 3.00 µm (Fig. 1c). Molds were fabricated from silicon-on-insulator wafers by electron-beam lithography and dry etching, and replicated in polydimethylsiloxane (PDMS) (Fig. 1d). Scanning electron microscopy (SEM) confirmed the formation of well-defined microchannels, with measured widths closely matching their designed dimensions (Fig. 1e). To confine bacterial cells, the PDMS device was simply placed onto a droplet of cell suspension on a glass surface, without the need for external flow control. Eight channel widths were arranged within a single microscopic field of view, allowing bacterial motility under different degrees of confinement to be observed simultaneously (Fig. 1c).

### Label-free tracking of cells in narrow channels

Phase-contrast imaging of bacteria in these channels is difficult because the strong contrast of the channel walls hides cells near the walls (Fig. 2a). To separate moving cells from the static channel structure without fluorescent labeling, we used temporal background subtraction. An intensity projection of the whole movie keeps the static device but weakens the signal from moving cells (Fig. 2b). Subtracting this projected image from each frame suppressed the channel background and enhanced the contrast of moving bacterial cells (Fig. 2c). The extracted cell signals were then overlaid on the background image to visualize cell positions relative to the channel geometry (Fig. 2d and Movie S1). Using the processed image sequences, we reconstructed continuous trajectories of individual cells within microchannels of different widths (Fig. 2e). Background subtraction increased the number of cells that could be tracked continuously for 5 s in the same field of view from 2 to 71 (Fig. 2f, g). This label-free tracking approach enabled quantitative analysis of bacterial swimming speeds and directional reversals under spatial confinement.

**Figure 2:**
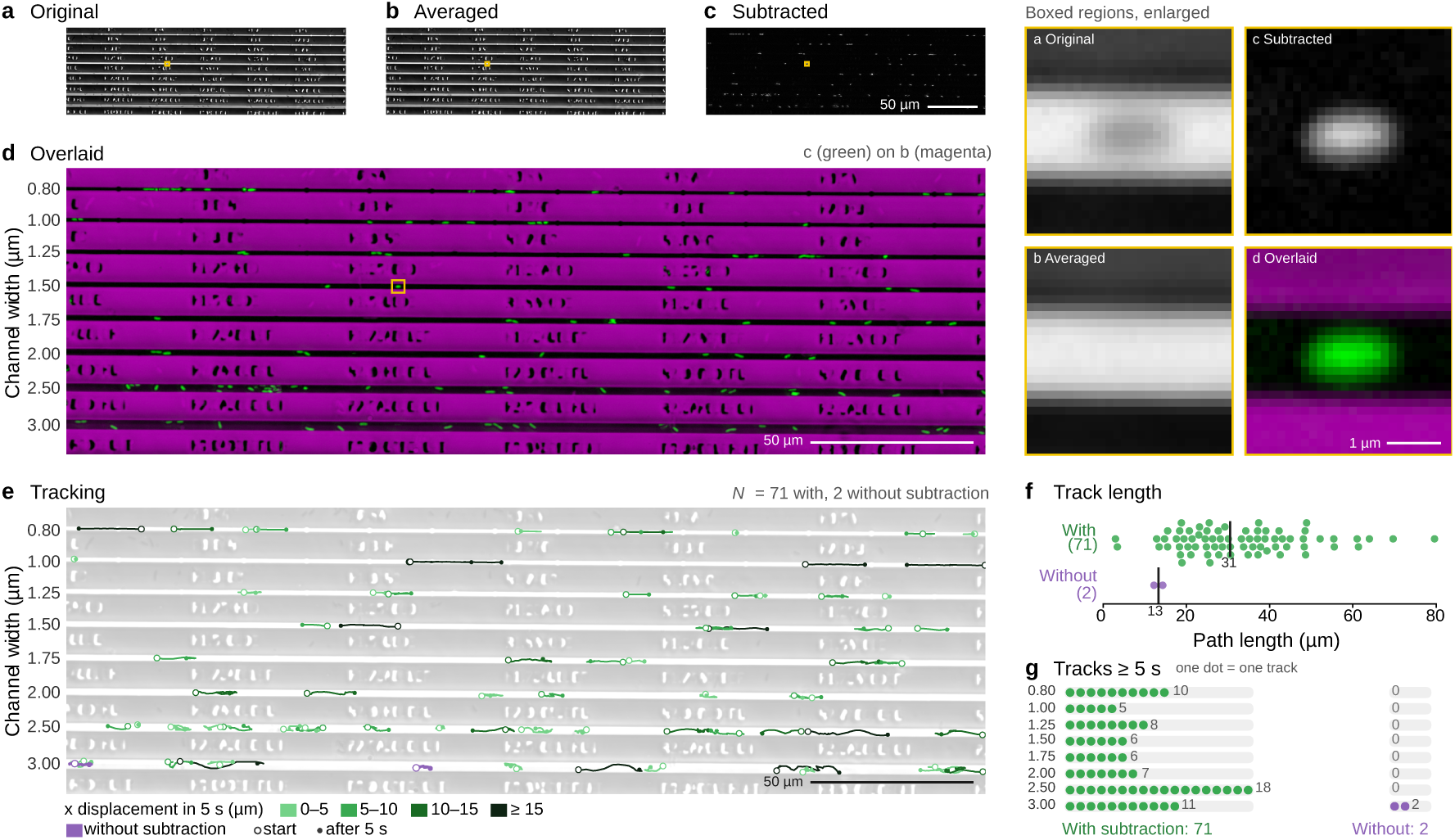
Label-free extraction and trajectory reconstruction of bacteria in cell-sized confinements. **a** Phase-contrast image of *C. insecticola* cells confined within microchannels. **b** Temporal intensity projection of the entire movie sequence. **c** Background-subtracted image obtained by subtracting the projected image from each frame of the original movie. **d** Pseudo-colored overlay of the projected image and the subtracted image. Channel structures and extracted bacterial signals are shown in magenta and green, respectively. The boxed regions in **a-d** are enlarged at the right. **e** Trajectories reconstructed from the processed image sequence. Cell trajectories recorded for 5 s at 0.033-s intervals are overlaid within the microchannels. **f** Path length over 5 s of each trajectory in **e**, with (green, *N* = 71) and without (purple, *N* = 2) background subtraction. Each dot represents one trajectory; vertical lines and numbers indicate the median path length (31 and 13 µm, respectively). **g** Number of trajectories lasting at least 5 s in each channel width, with (left, green) and without (right, purple) background subtraction. Each dot represents one trajectory, the number of trajectories is given next to each row, and the totals are given at the bottom.

### Physical environments change the speed ranking across species

We applied the device to eight species with five types of motility system (Fig. 3a). *C. insecticola* and *Burkholderia anthina* have a polar bundle of flagella, *V. fischeri* has a polar tuft of sheathed flagella, *Vibrio alginolyticus* has a single sheathed polar flagellum, *Salmonella enterica* and *Bacillus subtilis* have peritrichous flagella, *Campylobacter jejuni* has bipolar flagella, and *Flavobacterium johnsoniae* glides without flagella. We first measured each species in open chambers 5, 10 and 90 µm high, with or without 0.4% methylcellulose (MC), a polymer that is widely used to mimic the viscosity of mucus (Fig. 3b and Fig. S1). We then measured the same species in 2-µm-high channels containing MC (Fig. 3b, c, and Movie S2).

**Figure 3:**
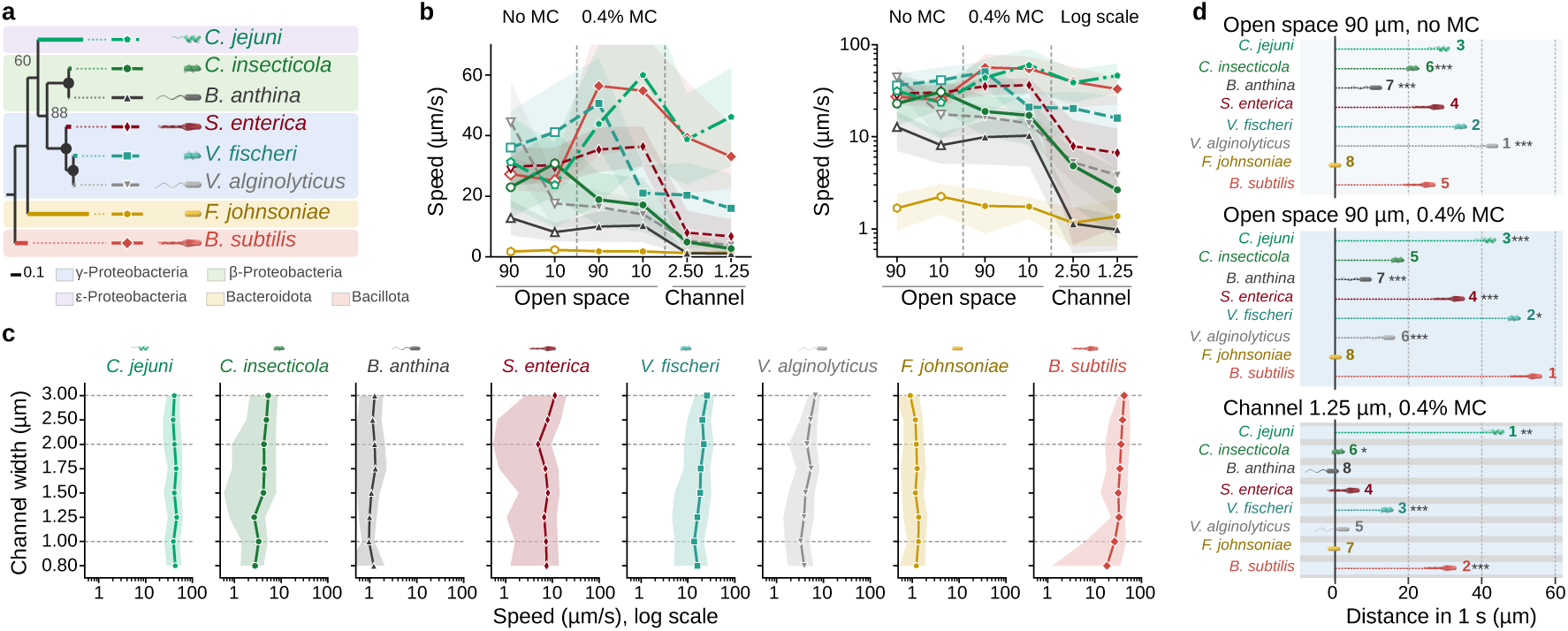
Physical environments reshape bacterial motility performance across species. **a** Phylogenetic relationships and motility systems of the eight bacterial species analyzed in this study. The tree was constructed from 16S rRNA gene sequences by the maximum-likelihood method; numbers at nodes are bootstrap values (%), and the scale bar indicates 0.1 substitutions per site. The species include the polar-flagellated bacteria *C. insecticola*, *B. anthina*, *V. fischeri*, and *V. alginolyticus*; the peritrichously flagellated bacteria *S. enterica* and *B. subtilis*; the bipolar-flagellated bacterium *C. jejuni*; and the gliding bacterium *F. johnsoniae*. Schematic illustrations indicate the characteristic motility mode and flagellar arrangement of each species. **b** Comparison of swimming speeds across representative physical environments. Mean motility speeds of the eight bacterial species were measured in conventional observation chambers under open-space conditions in the absence and the presence of 0.4% MC and in confined microchannels. Data are shown as mean ± SD (shaded areas). Left: Linear plot. Right: Logarithmic plot. Four open space and two narrow channel conditions including 90-µm height chamber without MC, 10-µm without MC, 90-µm with MC, 10-µm with MC, 2.50-µm-wide channel and 1.25-µm-wide channel with MC are presented. **c** Dependence of motility speed on microchannel width for the eight bacterial species. Each panel shows the mean swimming speed measured within microchannels of different widths (0.80–3.00 µm; channel height, 2 µm). Means and SD are presented. **d** Summary of rank changes in bacterial motility performance under different physical environments. Species are listed in the same order as in **a**, bars show the mean distance traveled in 1 s, numbers indicate the rank in each environment (1 = fastest; *N* = 20-167 cells per species and environment). In each environment, the speeds of species at adjacent ranks were compared (first versus second, second versus third, and so on). An asterisk next to a species indicates that it was significantly faster than the species ranked immediately below it (Welch’s t test with Holm correction, \**p* < 0.05, \*\**p* < 0.01, \*\*\**p* < 0.001).

Even in open chambers, the responses differed among species (Fig. 3d). In 90-µm chambers without MC, *V. alginolyticus* was the fastest of the eight species (44 ± 14 µm/s, mean ± SD). Reducing the chamber height to 10 µm lowered its speed to 18 ± 8 µm/s, and adding MC to the 90-µm chamber lowered it to 16 ± 5 µm/s. MC had the opposite effect on *B. subtilis* (27 ± 5 to 56 ± 23 µm/s), *C. jejuni* (31 ± 15 to 44 ± 26 µm/s) and *V. fischeri* (36 ± 12 to 51 ± 15 µm/s). The gliding bacterium *F. johnsoniae* moved at 1–2 µm/s under all conditions.

Lateral confinement separated the species further (Fig. 3c). *C. jejuni* kept a speed of 39–46 µm/s at all widths, and *B. subtilis* slowed gradually from 42 µm/s in 3.00-µm channels to 18 µm/s in 0.80-µm channels. In contrast, *V. alginolyticus* moved at only 3–7 µm/s at all channel widths. *S. enterica* also became much slower than in open chambers at 5–11 µm/s, whereas *F. johnsoniae* kept its slow but nearly constant gliding at all widths. The two species with peritrichous flagella behaved very differently. *B. subtilis* was among the fastest and *S. enterica* among the slower species, so flagellar arrangement alone does not predict performance under confinement. In addition, in narrow channels, the flagellated *B. anthina* was no faster than *F. johnsoniae*, which has no flagella at all (1.0 ± 0.3 versus 1.4 ± 0.6 µm/s in 1.00-µm channels).

As a result, the ranking of the eight species changed substantially (Fig. 3d and Movie S3). *V. alginolyticus* was first in 90-µm chambers without MC but fifth in 1.25-µm channels with MC. *B. subtilis* rose from fifth to second and *C. jejuni* from third to first, whereas *S. enterica* kept its rank as fourth. The fastest species was *V. alginolyticus* in open liquid without MC, *B. subtilis* or *V. fischeri* (not significantly different) in open chambers with MC, and *C. jejuni* in the channels. Only *C. jejuni* and *V. fischeri* remained among the top three in all three environments. Speed measured under conventional conditions therefore did not predict speed under cell-scale confinement, and no single species was the fastest in all environments.

We next compared the speed of each species in open liquid with the fraction of that speed retained under confinement (Fig. S2). Taking the ratio of speed in 1.25-µm channels to speed in 90-µm chambers without MC, *C. jejuni* and *B. subtilis* kept all of their open-liquid speed and the gliding *F. johnsoniae* most of its slow speed, whereas *V. fischeri* kept half, *S. enterica* about a quarter, and *C. insecticola*, *V. alginolyticus* and *B. anthina* only about a tenth. The two numbers were not correlated across the eight species. *V. alginolyticus* was fast in open liquid but lost most of its speed in the channels, whereas *C. jejuni* and *B. subtilis* were fast in both, and *F. johnsoniae* was slow in both.

Cell width explains part of this pattern but not all of it (Fig. S3). *C. jejuni* is a thin helical cell about 0.5 µm wide, so even a 0.80-µm channel hardly constrains it laterally, whereas *S. enterica* and *B. subtilis* are about 0.80 µm wide. Yet *B. subtilis* kept about 18 µm/s even in 0.80-µm channels, whereas *S. enterica*, which has a similar width, slowed strongly at all widths, and *B. anthina* was no faster than the thinner gliding *F. johnsoniae*.

### Host symbionts move faster than their non-symbiotic relatives under confinement

Differences under confinement were clearest between closely related species (Fig. 4). We compared two pairs, each consisting of a host symbiont and a relative that does not form the corresponding symbiosis, namely *C. insecticola* versus *B. anthina* (Betaproteobacteria), and *V. fischeri* versus *V. alginolyticus* (Gammaproteobacteria). In both pairs, the symbiont moved a long distance along the channel within 5 s, whereas the non-symbiont moved much less (Fig. 4a). In 1.50-µm channels, *C. insecticola* was about four times faster than *B. anthina*, and *V. fischeri* was about four times faster than *V. alginolyticus* (Fig. 4b and Fig. 3b, c). The advantage of the symbiont was seen at all channel widths examined. The two pairs differed in open liquid. Without MC, *V. alginolyticus* was faster than *V. fischeri* (44 ± 14 versus 36 ± 12 µm/s), but adding MC alone reversed their order (16 ± 5 versus 51 ± 15 µm/s), and confinement widened the gap further. *C. insecticola* was about twice as fast as *B. anthina* in open liquid with or without MC (23 versus 13 µm/s and 19 versus 10 µm/s, respectively), and confinement widened the gap to about four-fold.

**Figure 4:**
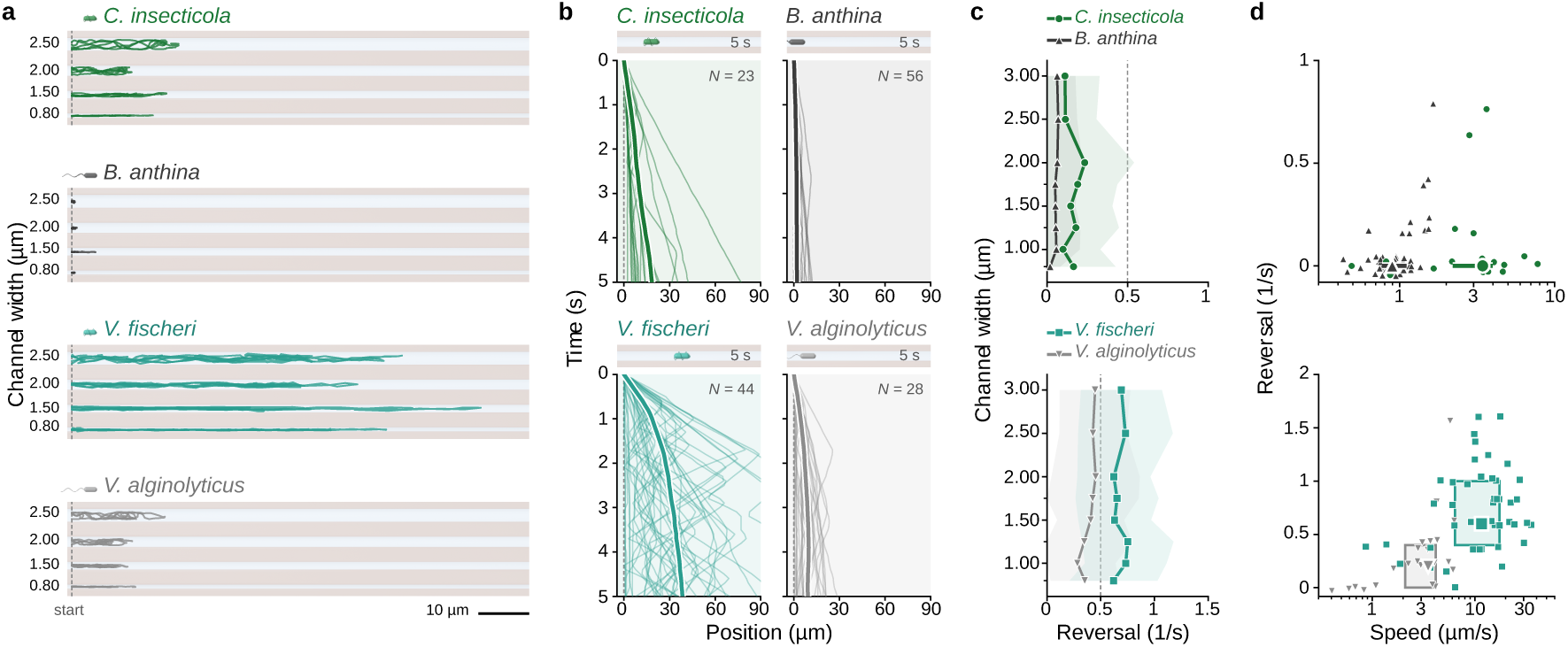
Distinct motility phenotypes among related bacteria under cell-scale confinement. **a** Trajectories of individual cells within microchannels of different widths. Five representative cell trajectories of *C. insecticola*, *B. anthina*, *V. fischeri*, and *V. alginolyticus* recorded over 5 s are shown for each species and channel width. The spacing between adjacent channels is schematic. **b** Time-dependent axial displacement of individual cells within 1.50-µm-wide microchannels. Thin lines represent trajectories of individual cells recorded over 5 s, and thick lines indicate their mean displacement. The numbers of analyzed cells are indicated in each panel. **c** Directional reversal frequency as a function of microchannel width. *Upper*: Comparison between *C. insecticola* and *B. anthina*. *Lower*: Comparison between *V. fischeri* and *V. alginolyticus*. Directional reversals were identified from changes in the sign of axial displacement over 0.3-s intervals, with a minimum displacement threshold of 0.5 µm. Lines indicate mean reversal frequencies, and shaded regions represent SD. **d** Relationship between swimming speed and directional reversal frequency for individual cells within 1.00-µm-wide microchannels. *Upper*: Comparison between *C. insecticola* and *B. anthina*. *Lower*: Comparison between *V. fischeri* and *V. alginolyticus*. Large symbols indicate median values, and rectangles represent the interquartile ranges of the two parameters.

The two symbionts moved in different ways. Plots of position along the channel against time showed that *C. insecticola* moved smoothly in one direction, whereas *V. fischeri* reversed often but still made rapid net progress (Fig. 4b). Reversal frequency, measured at each width, separated the two phylogenetic groups rather than the symbionts from the non-symbionts. Both betaproteobacteria reversed rarely (<0.25 s^−1^), and both *Vibrio* species reversed often (0.3–0.8 s^−1^), even though the speeds within each pair differed severalfold (Fig. 4c). Because a cell cannot turn around inside a channel narrower than its own length, a reversal detected in the channels corresponds to a change in the direction of propulsion, not to a reorientation of the cell. Reversal frequency therefore reflects how often propulsion switches direction independently of speed. When speed was plotted against reversal frequency for individual cells in 1.00-µm channels, the four species formed largely separate groups (Fig. 4d). *C. insecticola* was moderately fast with few reversals, *B. anthina* was slow with few reversals, *V. fischeri* fast with frequent reversals, and *V. alginolyticus* slow with frequent reversals. Thus, the two symbionts move efficiently through narrow channels in different ways.

### Both symbionts swim through narrow channels with wrapped flagella

To see how the symbionts keep moving in narrow channels, we imaged their flagella directly by fluorescence microscopy (Fig. 5). In *C. insecticola*, whose flagella were labeled with a Cy3 NHS ester, we classified each moving cell as unwrapped (flagella trailing behind the cell body) or wrapped (flagella wrapped around the cell body) (Fig. 5a and Movie S4). In channels 0.80–3.00 µm wide, the fraction of wrapping cells increased as the channels narrowed, from about 30% in 3.00-µm channels to about 50% in 1.00– 1.25-µm channels and 71% in 0.80-µm channels (Fig. 5b). Within each mode, speed changed little with channel width. Unwrapped cells moved at about 4–5 µm/s and wrapped cells at about 3–4 µm/s at all widths (Fig. 5c). Wrapped cells thus kept about 70–90% of the speed of unwrapped cells, a smaller penalty than in open liquid, where wrapped cells of this species swim at about one-third of the speed of unwrapped cells in buffer and about two-thirds in 0.3% MC^32^.

**Figure 5:**
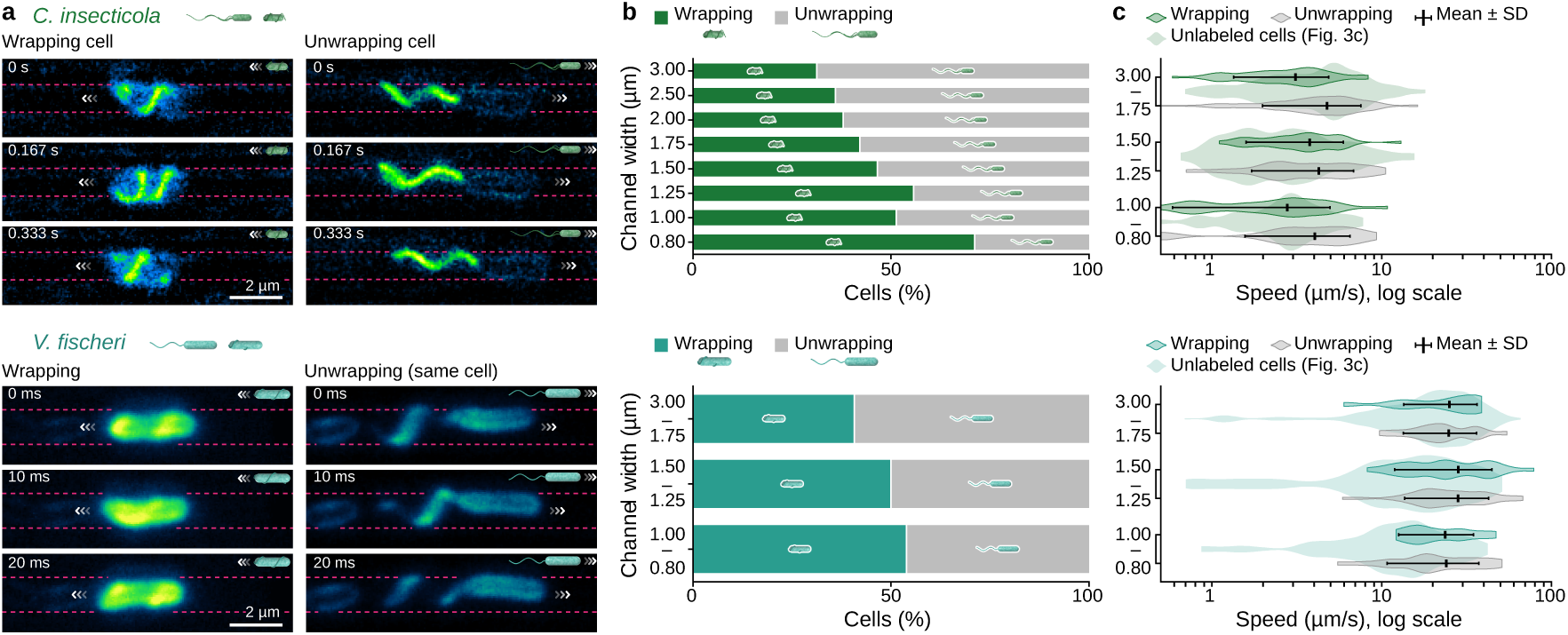
Flagellar wrapping of *C. insecticola* and *V. fischeri* in narrow microchannels. **a** Epifluorescence images of *C. insecticola* cells moving through a 1.00-µm-wide microchannel and TIRF images of *V. fischeri* cells moving through a 1.25-µm-wide microchannel in the unwrapping mode and the wrapping mode. Flagellar filaments and the cell surface of *C. insecticola* were labeled with Cy3 NHS ester. The sheathed polar flagella and the cell body of *V. fischeri* were stained with FM 4-64. The channel walls are shown with dashed lines. All images are shown without mirror reversal. In both species, the wrapped filament forms a left-handed helix. Scale bars, 2 µm. **b** Fraction of cells in the wrapping and unwrapping modes at each channel width. **c** Speed of individual cells in the wrapping and unwrapping modes in each group of channel widths. Violin plots show the distributions of individual-cell speeds, bars indicate the mean ± SD, and pale violins show unlabeled cells from Fig. 3c. The numbers of cells with measured speed are indicated below. All observations were made in medium supplemented with 0.4% (wt/vol) MC.

The polar flagella of *V. fischeri* are covered by a membrane sheath, so we stained them with the lipophilic dye FM 4-64 and recorded cells inside the channels (Fig. 5a and Movie S5). *V. fischeri* also passed through the channels with its flagella wrapped around the cell body, the wrapped mode previously described for this species in open liquid^31,32^. About 40–55% of the cells were wrapped in all groups of channel widths (Fig. 5b), and wrapped and unwrapped cells moved at similar speeds (about 20–30 µm/s, Fig. 5c). The wrapped filament formed a left-handed helix around the cell body in both species, with a pitch angle of 64 ± 11° in *C. insecticola* and 57 ± 9° in *V. fischeri* (Fig. S4), the same handedness as reported for wrapped filaments of *C. insecticola*, *Shewanella putrefaciens*, and *Pseudomonas putida* in open liquid^32,36,37^. The number of cells at some widths is small, but the observation shows that the wrapped mode is used under confinement by a symbiont from a second lineage, not only by *C. insecticola*^25^. Thus, in both pairs, the symbiont that kept moving under confinement swam with wrapped flagella, and inside the channels wrapping cost little or no speed.

## Discussion

Our results show that there is no universal champion of bacterial motility. The fastest species changed with every type of environmental change we applied, and some species moved up or down by as many as five ranks (Fig. 3). Motility performance is therefore a property of the organism and its environment together^42,43^, in the same way that a race time belongs to a runner on a particular track. One consequence is that swimming speeds measured in open liquid should not be extrapolated to the structured habitats in which many bacteria live. A second is that speed in open liquid and the fraction of speed retained under confinement were uncorrelated (Fig. S2). *V. alginolyticus* was fast but sensitive to confinement, whereas *F. johnsoniae* was slow but insensitive. Whether the two trade off against each other, or simply reflect different designs of the motility machinery, cannot be decided from eight species.

Adding MC alone produced the largest reordering, moving *B. subtilis* from fifth to first and *V. alginolyticus* from first to sixth, approximately doubling the speed of *B. subtilis* while reducing that of *V. alginolyticus* to about one-third (Fig. 3b and Fig. S2c). Enhanced swimming in polymer solutions has also been reported for *E. coli*^15,44,45^, potentially involving shear thinning, polymer depletion, and flagellar bundle stabilization ^45,46^. *B. subtilis* carries about 20–30 flagella per cell^47^, several times more than *S. enterica*, which may amplify such effects, although why *B. subtilis* gains so much remains unclear. The sodium-driven polar flagellum of *V. alginolyticus*, in contrast, is sensitive to viscosity and to nearby surfaces^48^, consistent with its loss of speed in shallow chambers even without MC. Lateral confinement imposed an additional constraint (Fig. 3c). It had little effect on *C. jejuni*, a thin helical cell that is known to keep swimming in viscous environments^49,50^, but a strong effect on *S. enterica*. *Salmonella* reaches the intestinal epithelium by swimming near surfaces and through gaps in mucus^51^, and its motility may be tuned to these conditions rather than to cell-sized passages. The gliding bacterium *F. johnsoniae* kept its slow speed at all widths. Gliding is driven by adhesins that move along the cell surface on a rotary motor^52,53^, and it requires contact with a surface, so a channel that touches the cell on several sides may be close to its normal working condition rather than an obstacle.

The difference was clearest between the two pairs of closely related species. In two distinct lineages, the host symbiont maintained a severalfold higher speed than its non-symbiotic relative at all channel widths examined (Fig. 4). The results are consistent with the idea that the narrow passages of their hosts act as filters on motility^25,42^. Flagellar wrapping has been proposed to help both symbionts move through the mucus-filled ducts of their hosts^32,33^. Two caveats apply. First, both symbionts also exhibit flagellar wrapping. It is difficult to distinguish the contributions of host association and wrapping ability. Second, host association by itself does not explain robustness to confinement. *S. enterica* slowed sharply, whereas *C. jejuni* maintained its speed. Thus, we propose that passage through cell-scale constrictions, rather than host association itself, favors confinement-robust motility. A recent finding in the bean bug is consistent with this view. A pathogenic *Burkholderia* strain related to the symbiont enters the same symbiotic organ through the same constriction and also shows flagellar wrapping^54^. More pairs of symbionts, pathogens and free-living relatives are needed to test this idea, and our device offers a practical assay.

The two symbionts kept moving in different ways, with persistent runs in *C. insecticola* and frequent reversals in *V. fischeri*, and reversal frequency followed phylogeny rather than symbiotic status (Fig. 4c). *V. alginolyticus* in open liquid alternates forward and backward runs and reorients by a flick that relies on a buckling instability of the flagellar hook^55^. Inside a channel narrower than the cell length, a flick cannot reorient the cell, so the same motor switching appears as back-and-forth movement along the channel axis. In channels, *V. alginolyticus* reversed frequently but made little net progress. In viscous open liquid, *V. fischeri* alternates between pushing and wrapped swimming, and in the wrapped state the motors reverse together with a switch of the filament between left- and right-handed forms, which keeps the cell moving backward for several seconds^31^. In the channels, the wrapped filaments of *V. fischeri* were left-handed (Fig. 5a) and the cells reversed frequently (Fig. 4c), suggesting that they switched between pushing and wrapped swimming without these long backward runs. Because the two modes moved at similar speeds in the channels (Fig. 5c), this frequent switching cost *V. fischeri* little speed.

Both symbionts passed through the channels with wrapped flagella (Fig. 5), and in *C. insecticola* the fraction of wrapped cells increased as the channels narrowed, while the speed of each mode stayed nearly constant. In open liquid, wrapping generally slows swimming, and its role differs among species. Wrapped cells of *C. insecticola* swim at about one-third of the speed of unwrapped cells^32^, and those of *V. fischeri* at one-third to one-half of the speed of pushing cells^31,32^. In *Pseudomonas aeruginosa*, the wrapped filament is right-handed, wrapped cells swim at about 40% of the pushing speed, and the wrapped state lasts about 1 s and mainly randomizes the swimming direction^34^. In *S. putrefaciens*, the wrapped flagellum moves the cell efficiently only near a surface, which allows the cell to escape from traps^36^. In our channels, by contrast, wrapped cells moved at 70–90% of the speed of unwrapped cells in both symbionts (Fig. 5c). This difference suggests that wrapping becomes efficient when the wrapped helix can push against a structure. For *C. insecticola,* the speed of wrapped cells relative to unwrapped cells was 0.25 in 15% Ficoll but 0.63 in 0.3% MC of similar viscosity, because MC slowed the rotation of extended flagella more than that of wrapped flagella and made each rotation of the wrapped flagellum about twice as effective as in Ficoll^32^. Near a surface, the wrapped flagellum moves the cell like a screw through a thread^32,33,36^, and hydrodynamic simulations predict that confinement barely slows a cell with a wrapped flagellum but slows a cell with an extended one^25,38^. A polymer network and nearby walls may therefore both act as a thread for the wrapped helix. Our channels contained MC and kept every cell close to a wall, which could explain why wrapped cells moved nearly as fast as unwrapped cells at all widths. Thus, in addition to turning, extending backward runs and escaping from traps in open liquid ^31,34,36^, wrapping can serve as an efficient mode of propulsion in cell-sized passages. Wrapping has also been reported in *C. jejuni*^16,49^, another species that maintained high swimming speeds under confinement, and in this bipolar species the flagellum at the leading pole wraps during normal fast swimming. Wrapping is not the only way to keep moving in a narrow channel. *B. subtilis* also retained most of its speed (Fig. 3). This species is a specialist of surfaces and interfaces that spreads over surfaces by swarming and by surfing on waves of surfactant^56^, and in soil it travels along fungal hyphae, the so-called fungal highway^57^. Whether these traits help a cell move in a PDMS channel is unknown.

Our approach has both advantages and clear limitations. Recording all widths in one field of view removes variation between field of view, illumination and samples, and background subtraction recovers trajectories without labels. These features set it apart from earlier multi-species designs, which used channels mostly wider than the cell and focused on trajectory patterns rather than performance^24^, and from our earlier screen in channels of a single width^25^. On the other hand, straight channels are a simplification of natural geometries, MC does not reproduce the full rheology of mucus, and short-time speed and reversal frequency do not describe long-distance migration or the probability of entering a constriction. Each species was measured in its own medium at 24 °C, more than 10 °C below the growth temperature of three of the species, and the channels were tested only with MC, so the effects of the polymer and of confinement cannot be fully separated in the channel data. Species that rarely entered the narrowest channels are represented only by the cells that did. The device measures components of motility performance, not ecological fitness.

In the Microbial Olympics, the winner depends on the event. Extending the comparison to more species, geometries and fluids should show how each motility system fits the structured habitats in which microbes actually compete.

## Materials and methods

### Device design and fabrication

Microfluidic devices were designed to examine bacterial motility under spatial confinement at dimensions comparable to individual bacterial cells (Movie S6). The devices consisted of parallel microchannels with a fixed height of 2.0 µm and designed widths of 0.80, 1.00, 1.25, 1.50, 1.75, 2.00, 2.50, and 3.00 µm (Fig. 1c). Eight channel widths ranging from 0.80 to 3.00 µm were used for quantitative motility comparisons.

Silicon molds were fabricated using silicon-on-insulator (SOI) wafers and electron-beam lithography. SOI wafers with a 2-µm-thick top silicon layer were diced into approximately 1-inch squares and spin-coated with a positive-tone electron-beam resist (ZEP-520A; Zeon, Japan). Channel patterns were defined by electron-beam lithography (F5112 + VD01; ADVANTEST, Japan), followed by deep reactive ion etching (DRIE) using a plasma etcher (MUC-21; Sumitomo Precision, Japan). The top silicon layer was etched down to the buried SiO₂ layer, allowing the thickness of the top silicon layer to define the microchannel height. The remaining electron-beam resist was removed by O₂ plasma ashing.

The silicon molds were treated with C₄F₈ plasma to form a hydrophobic release layer. Polydimethylsiloxane (PDMS; Sylgard 184; Dow, USA) was prepared by mixing the base polymer and curing agent at a ratio of 10:1 (w/w). The mixture was degassed under vacuum, poured onto the silicon mold, and subjected to a second vacuum degassing step to remove trapped air bubbles. The PDMS was cured at 110 °C for 20–30 min and carefully peeled from the mold. The resulting PDMS devices contained arrays of parallel microchannels with different widths.

### Characterization of microchannel geometry

The dimensions of the fabricated microchannels were evaluated by scanning electron microscopy (SEM). Channel dimensions were measured from calibrated SEM images using ImageJ/Fiji.

### Bacterial strains and culture conditions

Eight bacterial species representing different motility mechanisms were used in this study (Table S1). *C. insecticola* (formerly *Burkholderia insecticola*) RPE64 (JCM 31142) and *B. anthina* DSM 16086 cells were grown to an early log phase in 1/4 yeast-glucose (YG) liquid medium [0.125% (wt/vol) yeast extract, 0.100% (wt/vol) glucose and 0.025% (wt/vol) NaCl] at 28°C^40^. *V. fischeri* (*Aliivibrio fischeri*) ATCC 7744 cells were grown to an early log phase in seawater tryptone (SWT) liquid medium [0.5% (wt/vol) tryptone, 0.3% (wt/vol) yeast extract, 0.3% (wt/vol) glycerol, 70% (vol/vol) artificial seawater] with gentle shaking at 28°C^32^. *V. alginolyticus* Tj91 cells were grown in Marine broth (BD, USA) with shaking at 28°C^58^. *S. enterica* serovar Typhimurium strain SJW1103 and *B. subtilis* ATCC 6051 cells were grown in LB liquid medium [1% (wt/vol) tryptone, 0.5% (wt/vol) yeast extract and 1% (wt/vol) NaCl] with shaking at 37°C. *C. jejuni* EJC28 cells were cultured on Mueller-Hinton (MH) agar (Difco, USA) at 37°C under microaerobic conditions generated with a gas generator pack (AnaeroPack-MicroAero; Mitsubishi Gas Chemical, Japan)^49^. *F. johnsoniae* ATCC 17061 (UW101) cells were grown at 28°C in CYE medium [1% (wt/vol) polypeptone, 0.5% (wt/vol) yeast extract, 0.1% (wt/vol) MgSO_4_·7H_2_O, and 10 mM Tris-HCl (pH 7.5)], with shaking, to an optical density at 600 nm of 0.5 to 1.0^53^. The strains, motility characteristics, growth media, and cultivation temperatures are summarized in Table S1.

### Preparation of observation chambers and fluid conditions

Conventional observation chambers were constructed using two glass coverslips separated by parallel strips of double-sided adhesive tape. Three chamber heights were prepared using adhesive spacers with nominal thicknesses of 5 µm (8602TNW-05; DIC, Japan), 10 µm (7070W; Teraoka, Japan) and 90 µm (NW-5; Nichiban, Japan). To reduce the adhesion of large numbers of cells to the glass surface in the chambers, coverslips were glow-discharged using a hydrophilic treatment device (YHS-R; Sakigake-Semiconductor, Japan) for observations of *Vibrio* species, and chambers were pre-coated with 2% (wt/vol) BSA dissolved in 1/4 YG medium and washed with the same medium for observations of *B. anthina*. No surface treatment was applied for the other species. Bacterial suspensions were introduced between the coverslips, and the chambers were assembled to allow microscopic observation of cell movement. The chamber edges were sealed with nail polish. Bacterial motility was measured in the presence or absence of 0.4% (wt/vol) methylcellulose (MC4000, product M0512; Sigma-Aldrich, USA). Each bacterial species was suspended in its corresponding fresh observation medium, with or without MC. Thus, the base medium composition was maintained for comparisons between MC-supplemented and MC-free conditions within each species. Swimming speeds were measured in chambers with heights of 5, 10 and 90 µm under both fluid conditions, with the focal plane set near the surface of the lower coverslip.

### Confinement of bacteria in microfluidic devices

Bacterial cells were collected by centrifugation at 8,000 × *g* for 2 min and resuspended in fresh growth medium (Table S1) supplemented with 0.4% (wt/vol) methylcellulose (MC4000; Sigma-Aldrich, USA). For observations in the microfluidic device, the PDMS device was placed with the microchannel structures facing upward, and 6 µL of bacterial suspension was applied to the device. The device was then gently placed over a coverslip surface, enclosing bacterial cells within the microchannels. The assembled device was inverted for microscopic observation, and gentle pressure was applied to ensure contact between the PDMS and glass while avoiding air bubbles. Bacterial movement was observed within 20 min after assembly of the confinement device.

### Optical microscopy

Bacterial movements were visualized under a phase-contrast microscope (IX73; Olympus/Evident, Japan) equipped with 20×, 40×, or 100× objective lenses (UCPLFLN20×, LUCPLFLN40×, or UPLXAPO100×; Olympus/Evident, Japan), a CMOS camera (DMK 33UX174; Imaging Source, Germany), and an optical table (HAX-0806; JVI, Japan). Phase-contrast images were acquired as grayscale images at 10–80 fps and converted into an AVI file without any compression. Fluorescently labeled cells (Fig. 5, Fig. S4, Movie S4 and Movie S5) were observed with a fluorescence microscope equipped with a 100× objective lens (UPLanXAPO100×OPH, Olympus/Evident, Japan). For the *C. insecticola* cells in Fig. 5 and Movie S4, epifluorescence illumination was provided by a mercury lamp (U-HGLGPS, Olympus/Evident, Japan) through a filter set (Avant Cy3, Semrock, USA). All other fluorescence images, including the *V. fischeri* cells in Fig. 5 and Movie S5 and the cells of both species used for the pitch-angle measurements (Fig. S4), were recorded under a custom-built total internal reflection fluorescence (TIRF) microscopy, with a 532-nm laser (OBIS 532; Coherent, USA) through filters (Di02-R532; Semrock, USA, as the dichroic mirror and FELH550; Thorlabs, USA, as the emission filter) for *C. insecticola*, or with a 561-nm laser (OBIS 561; Coherent, USA) through filters (Di03-R488/561 as the dichroic mirror, and BLP01-568R as the emission filter; Semrock, USA) for *V. fischeri*. Images were recorded with CMOS cameras (DMK 33UX174; Imaging Source, Germany, and Zyla 4.2; Andor, UK). Fluorescence images were acquired as grayscale images at 30-200 fps and converted into a sequential TIF file without any compression. All data were analyzed by ImageJ 1.53k (rsb.info.nih.gov/ij/)^59^ and its plugin, TrackMate^60^. Observations were conducted at a room temperature of 24 °C.

### Image processing

Image sequences were processed using ImageJ (rsb.info.nih.gov/ij/)^59^. To generate a static representation of the microfluidic device, all frames in a movie were combined using the Z-projection function (maximum or median intensity), producing a projected image that primarily contained signals originating from the stationary channel structures. Because bacterial cells continuously moved throughout the recording, their contribution to the projected image was minimized. The absolute difference between each frame of the original movie and the projected image was then calculated using the Image Calculator function (Difference) in ImageJ, thereby removing signals associated with the microfluidic device and selectively retaining signals originating from moving bacterial cells. The processed image sequence was used for subsequent cell tracking and trajectory analysis. For visualization purposes, the extracted bacterial signals and the projected device image were occasionally displayed as pseudo-colored overlays, in which bacterial cells and channel structures were assigned distinct colors.

Image sequences recorded in open chambers were processed by the same background subtraction to improve the signal-to-noise ratio. Cells in the processed image sequences were tracked using TrackMate with the difference-of-Gaussian (DoG) detector and the simple linear assignment problem (LAP) tracker. For tracking without background subtraction (Fig. 2f), the original image sequences were inverted and tracked in the same manner. Erroneous tracks, such as those caused by tracking errors, cell overlap or ambiguous cell identification, were removed manually after tracking. All tracking data were subsampled to 100-ms intervals.

For each trajectory in the microchannels, the first continuous 5-s segment was analyzed. The speed of each cell was taken as the mean of 1-s path speeds, calculated with a 1-s window shifted by 100 ms along the 5-s segment. Trajectories whose maximum 1-s path speed was below 1 µm/s were excluded from the analysis. In open chambers, where cells were rarely tracked continuously for 5 s, the speed was calculated from the first 1 s of each trajectory. For *C. jejuni*, whose small and fast cells were difficult to track continuously, speeds were calculated from the first 0.5 s of each trajectory in both open chambers and microchannels. Directional reversals were detected from displacements along the channel axis over 0.3-s intervals. Only displacements of at least 0.5 µm were considered, and a reversal was counted when the sign of successive displacements changed. The reversal frequency was calculated as the number of reversals divided by the segment duration (5 s).

### Preparation of fluorescently labeled cells

Flagellar filaments of *C. insecticola* were labeled by a method slightly modified from the previous report^32^. Cell culture was centrifuged at 8,000 × g for 2 min at 24 °C and resuspended in phosphate-buffered saline (PBS) of 75 mM sodium phosphate (pH 7.4) and 68 mM NaCl supplemented with 20 mM glucose. The suspension was mixed with Sulfo-Cyanine3 NHS ester (Lumiprobe, USA) and incubated at 24 °C for 13 min. The suspension was washed three times in 1/4 YG liquid medium and resuspended in fresh 1/4 YG liquid medium supplemented with 0.4% (wt/vol) methylcellulose.

The polar flagella of *V. fischeri* are sheathed by the outer membrane and were stained with the lipophilic membrane dye FM 4-64 (Thermo Fisher Scientific, USA). Cells from an early-log-phase culture were collected by centrifugation at 8,000 × g for 2 min and resuspended in fresh SWT medium with 0.4% (wt/vol) MC containing 16 µM FM 4-64. The stained cells were confined in the microfluidic device as described above and imaged by TIRF microscopy at 5-ms intervals. Each moving cell was classified as wrapped or unwrapped from the position of the filament relative to the cell body, and its speed was measured from the same recording.

### Statistical analysis

Statistical analyses were performed with R (version 4.6.1). Speeds are given as the mean ± SD of individual cells. In each environment, the speeds of species at adjacent ranks were compared with two-sided Welch’s t tests (Fig. 3d and Movie S3), as were the speeds of each species in different conditions (Fig. S2), and *p* values were adjusted with the Holm method.

### Illustrations

Illustrations of bacterial cells, the bean bug and the bobtail squid, and schematics of the microfluidic device (Fig. 1, Fig. 3a and Movies S3–S6) were created by the authors in Blender (version 5.2; Blender Foundation, https://www.blender.org). Movies S3 and S6 were composed in Remotion (version 4.0.538; https://www.remotion.dev), a framework that renders videos from React code, from the tracking data, Blender renderings and, in Movie S6, the recording of Movie S1. No generative AI tool was used to create or modify any illustration, image or movie.

## Data and code availability

All data and code are available from the corresponding authors upon reasonable request.

## Acknowledgments

We thank Dr. Shoko Ueki for providing *V. alginolyticus* Tj91. This work was supported by JSPS KAKENHI Grant 24KJ1131 to NAU, 22H05066 and 26K01664 to DN, MEXT SPReAD program 26261082 to DN, JST FOREST Program JPMJFR2411 to DN, and MEXT ARIM program JPMXP12 25UT1130 to TK, the Samco Foundation to DN, the Naito Foundation to DN, and the Sasakawa Scientific Research Grant from The Japan Science Society (2025-6048) to YYS. The photolithography masks were made using the University of Tokyo Systems Design Lab (d.lab)’s 8-in. EB writer F5112 + VD01 donated by ADVANTEST Corporation. Microfabrication was performed in a clean room of the Division of Advanced Research Facilities (DARF) of the Coordinated Center for UEC Research Facilities of the University of Electro-Communications, Tokyo, Japan.

## Author contributions

Conceptualization: YYS, DN, and TK; Methodology: YYS, AY, NAU, and DN; Formal analysis and investigation: YYS, NAU, and DN; Funding acquisition: YYS, NAU, DN, and TK; Writing – original draft: YYS, NAU, DN, and TK; Writing – review & editing: DN.

## Competing interests

The authors declare that they have no competing interests.

## Supplementary Figures

### Movie Captions

**Movie S1: Simultaneous observation of *C. insecticola* motility in eight parallel microchannels of the microfluidic device.** Cells suspended in medium containing 0.4% MC were confined between the PDMS device and a glass coverslip, and then visualized under phase-contrast microscopy. Channel widths were 0.80, 1.00, 1.25, 1.50, 1.75, 2.00, 2.50 and 3.00 µm from top to bottom. Scale bar (upper right) and elapsed time in seconds (upper left) are shown. The movie has three parts, showing the original phase-contrast images, the processed images with the microfluidic device colored magenta and moving cells highlighted in green, and the processed images with the reconstructed trajectories in yellow. The movie is played in real time.

**Movie S2: Motility of the eight bacterial species in the parallel microchannels.** Cells of each species, suspended in their growth medium containing 0.4% MC, were confined between the PDMS device and a glass coverslip and recorded by phase-contrast microscopy. The recordings are shown after background subtraction, with the static channel structure pseudo-colored in magenta and moving cells in green (see Materials and Methods). The first half of the movie shows *C. jejuni*, *C. insecticola*, *B. anthina,* and *S. enterica*, and the second half shows *V. fischeri*, *V. alginolyticus*, *F. johnsoniae*, and *B. subtilis*. Channel widths are 0.80, 1.00, 1.25, 1.50, 1.75, 2.00, 2.50 and 3.00 µm from top to bottom. Scale bar and elapsed time in seconds are shown. The movie is played in real time.

**Movie S3: Comparison of the swimming speeds of the eight species in three conditions.** The eight species are shown as a race in (I) open 90-µm-high chambers without MC, (II) open 90-µm-high chambers with 0.4% MC and (III) 1.25-µm-wide microchannels with 0.4% MC. Each species is represented by a cell icon that advances along a representative trajectory reconstructed from the single-cell tracking data, and 1 s of swimming is shown over 4 s. The horizontal axis gives the distance from the start. At the end of each race, the rank and mean speed of each species (mean of all cells; *N* = 20–167 per species and condition) are displayed. Arrows next to each rank indicate the change in rank from the previous race. In each environment, species at adjacent ranks were compared, and an asterisk next to a species indicates that it was significantly faster than the species ranked immediately below it (Welch’s *t* test with Holm correction; \**p* < 0.05, \*\**p* < 0.01, \*\*\**p* < 0.001). No mark indicates no significant difference. The final chart summarizes the changes in rank across the three environments and corresponds to Fig. 3d.

**Movie S4: Flagellar dynamics of *C. insecticola* swimming in a narrow microchannel.** Flagellar filaments and the cell surface were labeled with Cy3 NHS ester, and cells suspended in 1/4 YG medium containing 0.4% MC were confined in the microfluidic device and recorded by epifluorescence microscopy at 17-ms intervals in a 1.00-µm-wide channel. The movie shows a wrapping cell, whose flagellar bundle wraps around the cell body, and an unwrapping cell, whose bundle trails behind the cell body, as in Fig. 5a. The two cells were recorded separately and are shown first one after the other and then side by side. Fluorescence intensity is shown in pseudo-color, dashed lines mark the channel walls, and schematic cells indicate the swimming mode. The movie is played in real time.

**Movie S5: Flagellar dynamics of *V. fischeri* swimming in a narrow microchannel.** The sheathed polar flagella and the cell body were stained with the lipophilic membrane dye FM 4-64, and cells suspended in SWT medium containing 0.4% MC were confined in the microfluidic device and recorded by TIRF microscopy at 5-ms intervals in a 1.25-µm-wide channel. The movie shows a cell switching from the wrapping mode to the unwrapping mode, in which the flagellum trails behind the cell body, as in Fig. 5a. The 0.7-s recording of the same cell is played 10 times slower than real time, first in sequence and then with the wrapped and unwrapped phases side by side. Fluorescence intensity is shown in pseudo-color, dashed lines mark the channel walls, and schematic cells indicate the swimming mode.

**Movie S6: Fabrication of the microfluidic device and confinement of bacterial cells.** Animated overview of the fabrication process described in Methods and summarized in Fig. 1d. A silicon-on-insulator (SOI) wafer with a 2-µm-thick silicon device layer is spin-coated with a positive electron-beam resist, the channel pattern is written by electron-beam lithography and developed, and the device layer is etched down to the buried oxide by deep reactive ion etching, so that the thickness of the device layer defines the channel height. After removal of the resist by O_2_ plasma ashing and deposition of a hydrophobic C_4_F_8_ release layer, PDMS is poured onto the silicon mold, degassed under vacuum, cured at 110 °C for 20–30 min and peeled off. A droplet of bacterial suspension is then sandwiched between the PDMS device and a cover glass, which encloses the cells in the parallel microchannels of eight widths within a single field of view. An inset shows a schematic cross-section at each step. The final segment shows a phase-contrast recording of cells in the eight channels after background subtraction, with the device in magenta and moving cells in green (as in Movie S1).

---

**Table S1.** Strain details and culture conditions.

| Strain | Description | Medium | Temp (℃) | Reference |
|---|---|---|---|---|
| *Caballeronia insecticola* | Betaproteobacteria, lophotrichous flagellation, flagellar wrapping | 1/4 YG | 28 | JCM 31142 |
| *Burkholderia anthina* | Betaproteobacteria, lophotrichous flagellation | 1/4 YG | 28 | DSM 16086 |
| *Vibrio fischeri* | Gammaproteobacteria, lophotrichous flagellation, flagellar wrapping | SWT | 28 | ATCC 7744 |
| *Vibrio alginolyticus* | Gammaproteobacteria, monotrichous flagellation | Marine broth | 28 | Tj91<sup>58</sup> |
| *Salmonella enterica* | Gammaproteobacteria, peritrichous flagellation | LB | 37 | SJW1103 |
| *Bacillus subtilis* | Bacillota (Firmicutes), peritrichous flagellation | LB | 37 | ATCC 6051 |
| *Campylobacter jejuni* | Campylobacterota, amphitrichous flagellation, flagellar wrapping | MH | 37 | EJC28<sup>49</sup> |
| *Flavobacterium johnsoniae* | Bacteroidota, gliding without flagella | CYE | 28 | ATCC 17061 |

**Figure S1:**
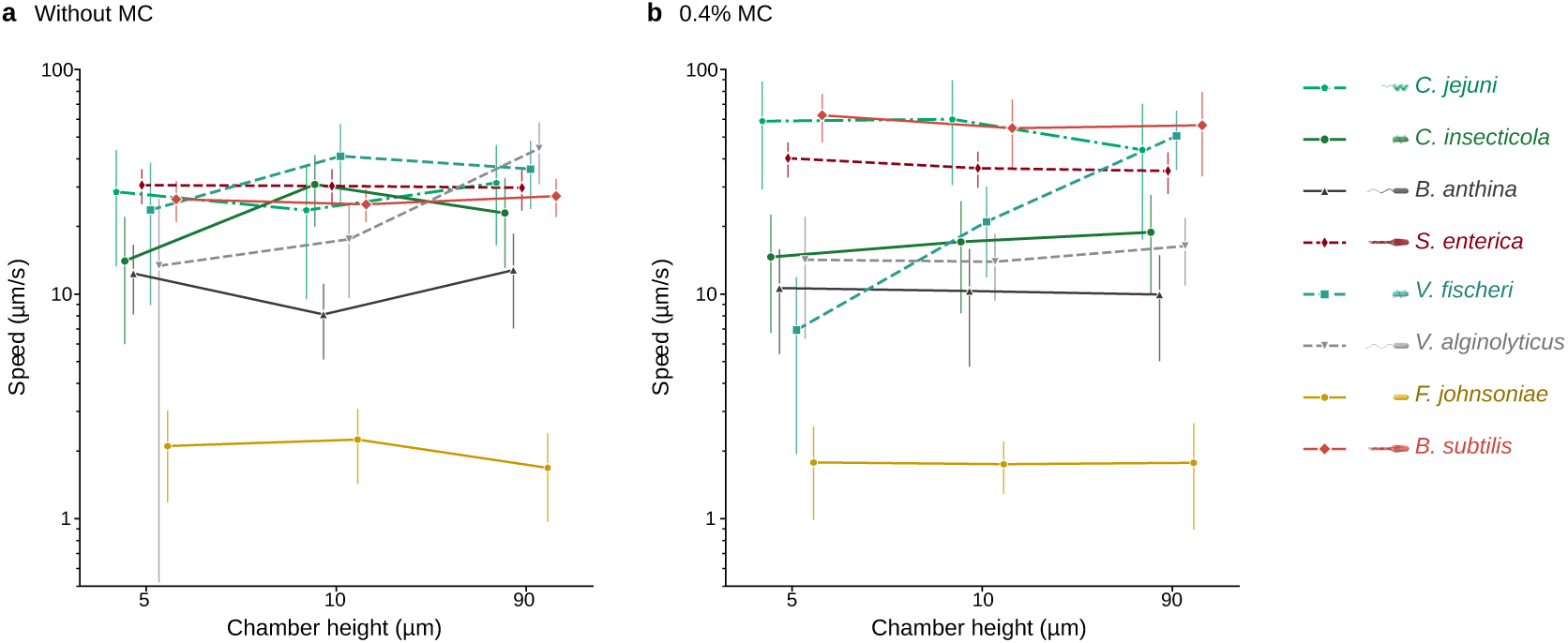
Effects of chamber height and methylcellulose supplementation on the motility speeds of eight bacterial species. **a** Swimming speeds in the absence of MC. **b** Swimming speeds in the presence of 0.4% MC. Mean swimming speed is plotted on a logarithmic scale against chamber height (5, 10 and 90 µm) for each of the eight bacterial species. Data are means ± SD from at least 20 individual cells per condition. For each cell, speed was calculated from the first 1 s of its trajectory (0.5 s for *C. jejuni*; see Materials and Methods). Symbols are offset slightly along the horizontal axis to avoid overlap. Species are identified by the colors and symbols in the key on the right, which are the same as in Fig. 3; the data for 90-µm chambers are those shown in Fig. 3b.

**Figure S2:**
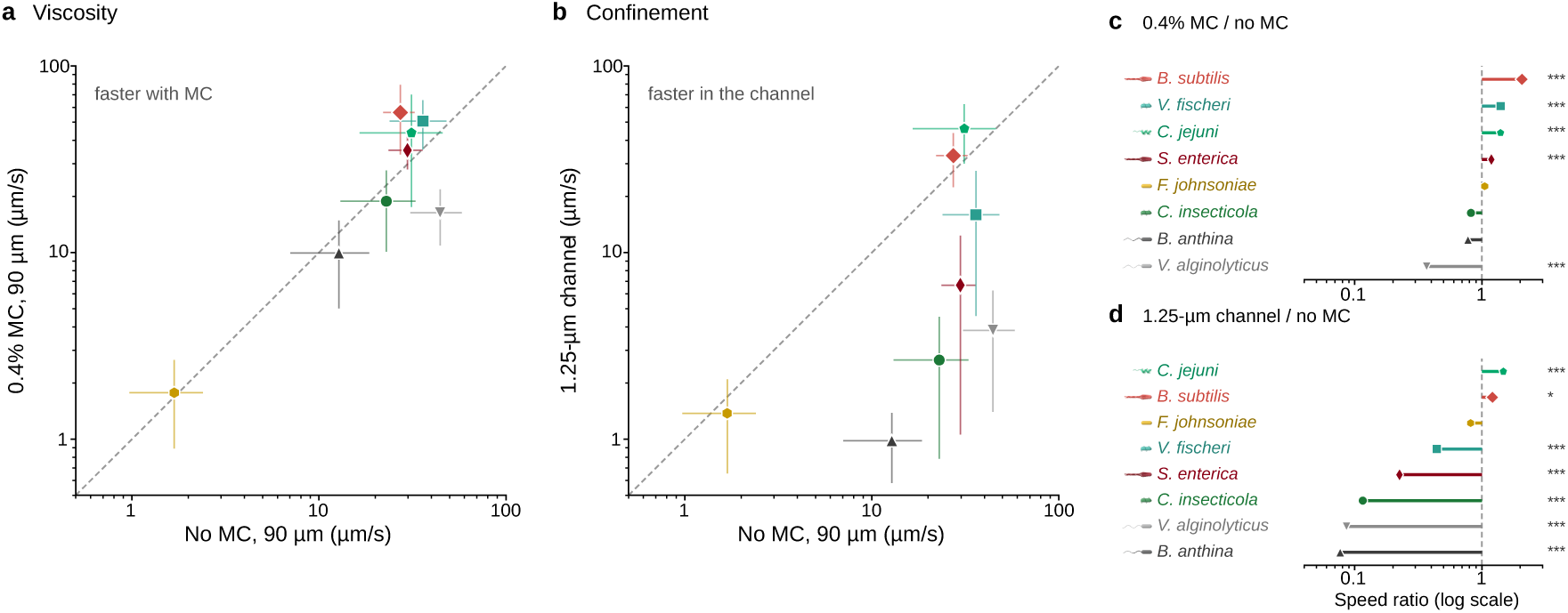
Effects of methylcellulose and cell-scale confinement on the swimming speeds of eight bacterial species. **a** Effect of viscosity. Mean speed in 90-µm-high chambers with 0.4% MC is plotted against mean speed in 90-µm-high chambers without MC. **b** Effect of confinement. Mean speed in 1.25-µm-wide microchannels with 0.4% MC is plotted against mean speed in 90-µm-high chambers without MC. In **a** and **b**, both axes are logarithmic, error bars indicate SD, and the dashed diagonal line marks equal speed in the two conditions. Species above the line were faster with MC (**a**) or in the channel (**b**). **c** Speed ratio of 0.4% MC to no MC in 90-µm-high chambers, calculated from the mean speeds in **a**. **d** Speed ratio of 1.25-µm-wide channels with MC to 90-µm-high chambers without MC, calculated from the mean speeds in **b**. In **c** and **d**, species are ordered by the ratio, the horizontal axis is logarithmic, and values above 1 indicate faster and values below 1 slower movement than in the reference condition. Mean speeds and SD were calculated from at least 20 individual cells per condition. Asterisks indicate statistically significant differences in individual-cell speeds between the test and the corresponding reference conditions (Welch’s t test with Holm correction; \**p* < 0.05, \*\**p* < 0.01, \*\*\**p* < 0.001). Species are identified by the colors and symbols used in Fig. 3.

**Figure S3:**
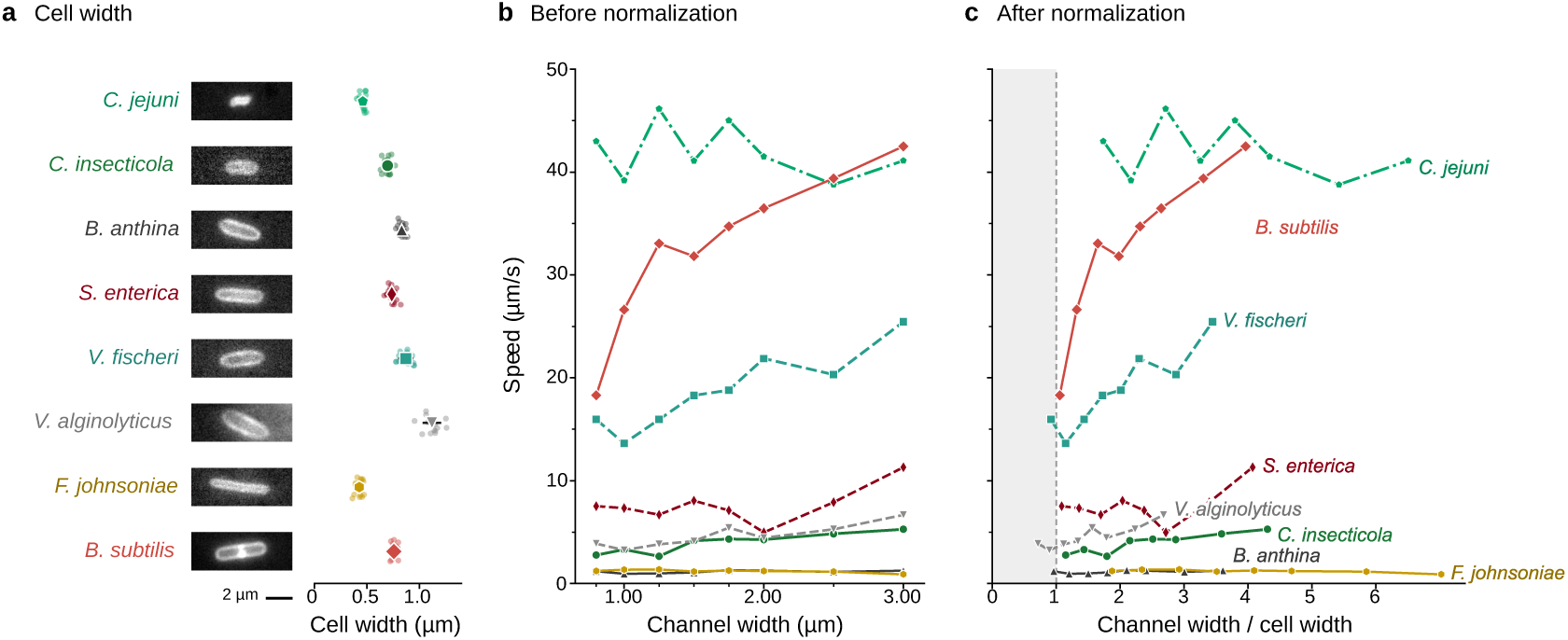
Cell width and swimming speed under cell-scale confinement. **a** Cell widths of the eight bacterial species. Left: Fluorescence images of representative cells stained with the membrane dye FM 4-64 of the eight species used in this study. Scale bar, 2 µm. Right: Cell widths measured from the fluorescence images. Dots represent 15 individual cells per species, and bars indicate the mean ± SD. **b** Mean swimming speed plotted against channel width (0.80–3.00 µm; channel height, 2 µm), as in Fig. 3c, with speed on a logarithmic scale. **c** The same data plotted against channel width normalized by the mean cell width of each species (channel width/cell width). A value of 1 on the horizontal axis indicates a channel as wide as the cell, and the gray area marks channels narrower than the cell. The eight bacterial species are distinguished by colors and labels.

**Figure S4:**
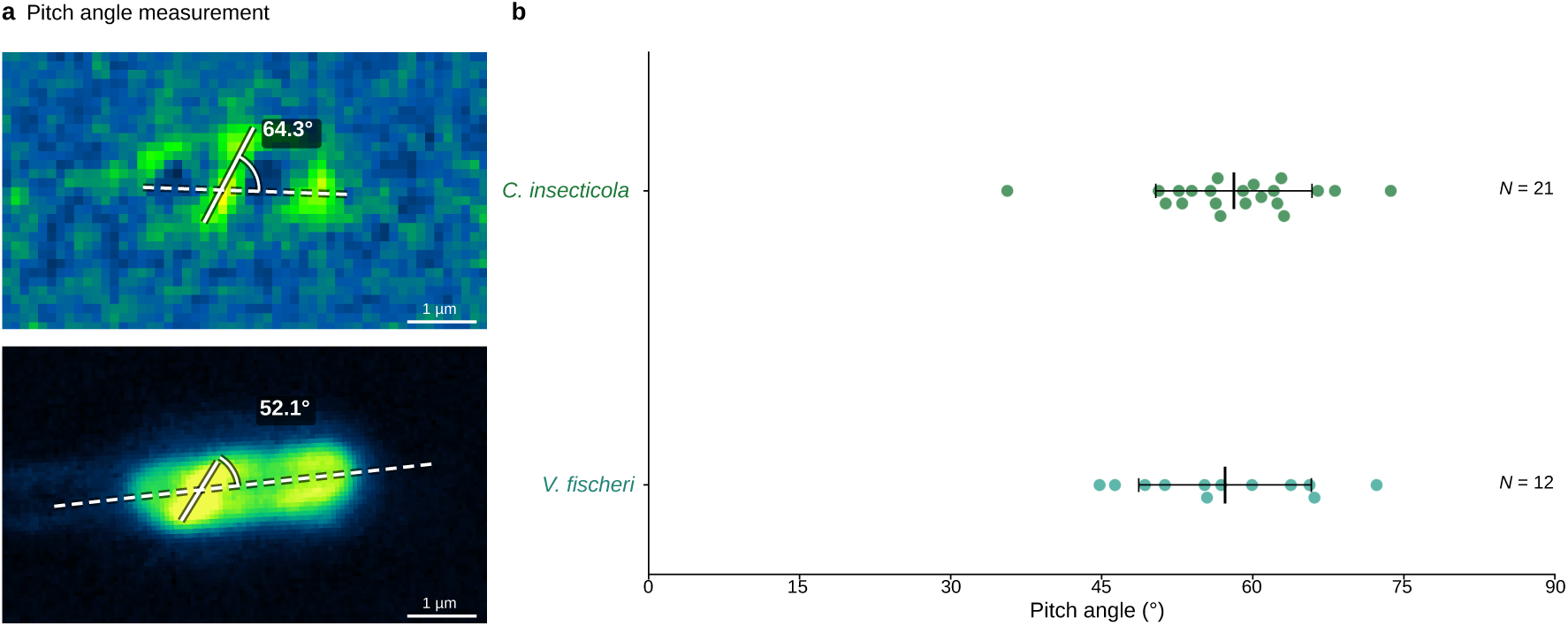
Helical pitch angle of wrapped flagellar filaments. **a** Measurement of the pitch angle on TIRF images of wrapped cells of *C. insecticola* (top) and *V. fischeri* (bottom). The dashed line marks the long axis of the cell, which is taken as the axis of the helix formed by the wrapped filament. The solid line follows the filament where it crosses the cell body. The pitch angle was measured as the angle between the two lines, and the measured value is shown for each cell. Flagella were labeled with Cy3 NHS ester (*C. insecticola*) or stained with FM 4-64 (*V. fischeri*) as in Fig. 5, and fluorescence intensity is shown in pseudo-color. Scale bars, 1 µm. **b** Pitch angles of the wrapped filaments of *C. insecticola* (*N* = 21 cells) and *V. fischeri* (*N* = 12 cells). Dots represent individual cells, and bars indicate the mean ± SD. The angle was measured counterclockwise from the cell axis to the filament. Because TIRF illumination shows only the part of the filament closest to the coverslip and the images are shown without mirror reversal, a counterclockwise tilt indicates a left-handed helix.

## Notes

### Competing Interest Statement

The authors have declared no competing interest.

## References

1 Wadhwa, N. & Berg, H. C. Bacterial motility: machinery and mechanisms. Nat Rev Microbiol 20, 161–173 (2022). 10.1038/s41579-021-00626-4

2 Keegstra, J. M., Carrara, F. & Stocker, R. The ecological roles of bacterial chemotaxis. Nat Rev Microbiol 20, 491–504 (2022). 10.1038/s41579-022-00709-w

3 Nelson, M. B. et al. The microbial Olympics 2016. Nat Microbiol 1, 16122 (2016). 10.1038/nmicrobiol.2016.122

4 Youle, M. et al. The microbial Olympics. Nat Rev Microbiol 10, 583–588 (2012). 10.1038/nrmicro2837

5 Maiuri, P. et al. The first World Cell Race. Curr Biol 22, R673–675 (2012). 10.1016/j.cub.2012.07.052

6 Conrad, J. C. & Poling-Skutvik, R. Confined flow: Consequences and implications for bacteria and biofilms. Annu Rev Chem Biomol Eng 9, 175–200 (2018). 10.1146/annurev-chembioeng-060817-084006

7 Lauga, E. Bacterial hydrodynamics. Annu Rev Fluid Mech 48, 105–130 (2016). 10.1146/annurev-fluid-122414-034606

8 Nord, A. L. et al. Catch bond drives stator mechanosensitivity in the bacterial flagellar motor. Proc Natl Acad Sci U S A 114, 12952–12957 (2017). 10.1073/pnas.1716002114

9 Beeby, M. et al. Diverse high-torque bacterial flagellar motors assemble wider stator rings using a conserved protein scaffold. Proc Natl Acad Sci U S A 113, E1917–E1926 (2016). 10.1073/pnas.1518952113

10 DiLuzio, W. R., et al. *Escherichia coli* swim on the right-hand side. Nature 435, 1271–1274 (2005). 10.1038/nature03660

11 Männik, J., Driessen, R., Galajda, P., Keymer, J. E. & Dekker, C. Bacterial growth and motility in sub-micron constrictions. Proc Natl Acad Sci U S A 106, 14861–14866 (2009). 10.1073/pnas.0907542106

12 Vizsnyiczai, G. et al. A transition to stable one-dimensional swimming enhances *E. coli* motility through narrow channels. Nat Commun 11, 2340 (2020). 10.1038/s41467-020-15711-0

13 Zhu, L., Lauga, E. & Brandt, L. Low-Reynolds-number swimming in a capillary tube. J Fluid Mech 726, 285–311 (2013). 10.1017/jfm.2013.225

14 Spagnolie, S. E. & Underhill, P. T. Swimming in complex fluids. Annu Rev Condens Matter Phys 14, 381–415 (2023). 10.1146/annurev-conmatphys-040821-112149

15 Martinez, V. A. et al. Flagellated bacterial motility in polymer solutions. Proc Natl Acad Sci U S A 111, 17771–17776 (2014). 10.1073/pnas.1415460111

16 Grognot, M. & Taute, K. M. More than propellers: how flagella shape bacterial motility behaviors. Curr Opin Microbiol 61, 73–81 (2021). 10.1016/j.mib.2021.02.005

17 McBride, M. J. *Bacteroidetes* gliding motility and the type IX secretion system. Microbiol Spectr 7, 10.1128/microbiolspec.psib-0002-2018 (2019). 10.1128/microbiolspec.psib-0002-2018

18 Gurung, J. P., Gel, M. & Baker, M. A. B. Microfluidic techniques for separation of bacterial cells via taxis. Microb Cell 7, 66–79 (2020). 10.15698/mic2020.03.710

19 Rusconi, R., Garren, M. & Stocker, R. Microfluidics expanding the frontiers of microbial ecology. Annu Rev Biophys 43, 65–91 (2014). 10.1146/annurev-biophys-051013-022916

20 Hol, F. J. H. & Dekker, C. Zooming in to see the bigger picture: Microfluidic and nanofabrication tools to study bacteria. Science 346, 1251821 (2014). 10.1126/science.1251821

21 Lynch, J. B. et al. Transitioning to confined spaces impacts bacterial swimming and escape response. Biophys J 121, 2653–2662 (2022). 10.1016/j.bpj.2022.04.008

22 Kurzthaler, C. et al. A geometric criterion for the optimal spreading of active polymers in porous media. Nat Commun 12, 7088 (2021). 10.1038/s41467-021-26942-0

23 Bhattacharjee, T. & Datta, S. S. Bacterial hopping and trapping in porous media. Nat Commun 10, 2075 (2019). 10.1038/s41467-019-10115-1

24 Tokárová, V. et al. Patterns of bacterial motility in microfluidics-confining environments. Proc Natl Acad Sci U S A 118, e2013925118 (2021). 10.1073/pnas.2013925118

25 Yoshioka, A. et al. Bacteria break through one-micrometer-square passages by flagellar wrapping. Nat Commun 17, 713 (2026). 10.1038/s41467-025-67507-9

26 Mergaert, P., Jung, M., Kikuchi, Y. & Ohbayashi, T. Assembly by disassembly of the gut microbiota in *Riptortus pedestris* and other stinkbugs. Curr Opin Insect Sci 79, 101622 (2027). 10.1016/j.cois.2026.101622

27 Ohbayashi, T. et al. Insect’s intestinal organ for symbiont sorting. Proc Natl Acad Sci U S A 112, E5179–E5188 (2015). 10.1073/pnas.1511454112

28 Kikuchi, Y., Hosokawa, T. & Fukatsu, T. Insect-microbe mutualism without vertical transmission: a stinkbug acquires a beneficial gut symbiont from the environment every generation. Appl Environ Microbiol 73, 4308–4316 (2007). 10.1128/aem.00067-07

29 Visick, K. L., Stabb, E. V. & Ruby, E. G. A lasting symbiosis: how *Vibrio fischeri* finds a squid partner and persists within its natural host. Nat Rev Microbiol 19, 654–665 (2021). 10.1038/s41579-021-00557-0

30 Sycuro, L. K., Ruby, E. G. & McFall-Ngai, M. Confocal microscopy of the light organ crypts in juvenile *Euprymna scolopes* reveals their morphological complexity and dynamic function in symbiosis. J Morphol 267, 555–568 (2006). 10.1002/jmor.10422

31 Zhuang, X.-Y., Tseng, C.-K. & Lo, C.-J. Viscosity-dependent swimming patterns assist *Aliivibrio* fischeri for symbiont partner finding. Phys Rev Res 6, 033214 (2024). 10.1103/PhysRevResearch.6.033214

32 Kinosita, Y., Kikuchi, Y., Mikami, N., Nakane, D. & Nishizaka, T. Unforeseen swimming and gliding mode of an insect gut symbiont, Burkholderia sp. RPE64, with wrapping of the flagella around its cell body. ISME J 12, 838–848 (2018). 10.1038/s41396-017-0010-z

33 Thormann, K. M., Beta, C. & Kühn, M. J. Wrapped up: The motility of polarly flagellated bacteria. Annu Rev Microbiol 76, 349–367 (2022). 10.1146/annurev-micro-041122-101032

34 Tian, M., Wu, Z., Zhang, R. & Yuan, J. A new mode of swimming in singly flagellated *Pseudomonas aeruginosa*. Proc Natl Acad Sci U S A 119, e2120508119 (2022). 10.1073/pnas.2120508119

35 Constantino, M. A. et al. Bipolar lophotrichous *Helicobacter suis* combine extended and wrapped flagella bundles to exhibit multiple modes of motility. Sci Rep 8, 14415 (2018). 10.1038/s41598-018-32686-7

36 Kühn, M. J., Schmidt, F. K., Eckhardt, B. & Thormann, K. M. Bacteria exploit a polymorphic instability of the flagellar filament to escape from traps. Proc Natl Acad Sci U S A 114, 6340–6345 (2017). 10.1073/pnas.1701644114

37 Hintsche, M. et al. A polar bundle of flagella can drive bacterial swimming by pushing, pulling, or coiling around the cell body. Sci Rep 7, 16771 (2017). 10.1038/s41598-017-16428-9

38 Gidituri, H., Ellero, M. & Balboa Usabiaga, F. Swimming efficiently by wrapping. J Fluid Mech 993, A7 (2024). 10.1017/jfm.2024.594

39 Kataoka, T., Yoneda, T., Nakane, D. & Wada, H. Bacteria harness torque-induced buckling instability for flagellar wrapping. Phys Rev E 113, 054402 (2026). 10.1103/ktss-zp41

40 Itoh, H. et al. Host–symbiont specificity determined by microbe–microbe competition in an insect gut. Proc Natl Acad Sci U S A 116, 22673–22682 (2019). 10.1073/pnas.1912397116

41 Nyholm, S. V. & McFall-Ngai, M. J. Dominance of *Vibrio fischeri* in secreted mucus outside the light organ of Euprymna scolopes: the first site of symbiont specificity. Appl Environ Microbiol 69, 3932–3937 (2003). 10.1128/aem.69.7.3932-3937.2003

42 Raina, J.-B., Fernandez, V., Lambert, B., Stocker, R. & Seymour, J. R. The role of microbial motility and chemotaxis in symbiosis. Nat Rev Microbiol 17, 284–294 (2019). 10.1038/s41579-019-0182-9

43 Mitchell, J. G. & Kogure, K. Bacterial motility: links to the environment and a driving force for microbial physics. FEMS Microbiol Ecol 55, 3–16 (2006). 10.1111/j.1574-6941.2005.00003.x

44 Torres Maldonado, B. O., et al. Enhancement of bacterial rheotaxis in non-Newtonian fluids. Proc Natl Acad Sci U S A 121, e2417614121 (2024). 10.1073/pnas.2417614121

45 Zöttl, A. & Yeomans, J. M. Enhanced bacterial swimming speeds in macromolecular polymer solutions. Nat Phys 15, 554–558 (2019). 10.1038/s41567-019-0454-3

46 Qu, Z., Temel, F. Z., Henderikx, R. & Breuer, K. S. Changes in the flagellar bundling time account for variations in swimming behavior of flagellated bacteria in viscous media. Proc Natl Acad Sci U S A 115, 1707–1712 (2018). 10.1073/pnas.1714187115

47 Guttenplan, S. B., Shaw, S. & Kearns, D. B. The cell biology of peritrichous flagella in *Bacillus subtilis*. Mol Microbiol 87, 211–229 (2013). 10.1111/mmi.12103

48 Atsumi, T. et al. Effect of viscosity on swimming by the lateral and polar flagella of *Vibrio alginolyticus*. J Bacteriol 178, 5024–5026 (1996). 10.1128/jb.178.16.5024-5026.1996

49 Cohen, E. J., et al. *Campylobacter jejuni* motility integrates specialized cell shape, flagellar filament, and motor, to coordinate action of its opposed flagella. PLOS Pathog 16, e1008620 (2020). 10.1371/journal.ppat.1008620

50 Ferrero, R. L. & Lee, A. Motility of *Campylobacter jejuni* in a viscous environment: comparison with conventional rod-shaped bacteria. J Gen Microbiol 134, 53–59 (1988). 10.1099/00221287-134-1-53

51 Furter, M., Sellin, M. E., Hansson, G. C. & Hardt, W.-D. Mucus architecture and near-surface swimming affect distinct *Salmonella* Typhimurium infection patterns along the murine intestinal tract. Cell Rep 27, 2665–2678.e2663 (2019). 10.1016/j.celrep.2019.04.106

52 Shrivastava, A., Lele, P. P. & Berg, H. C. A rotary motor drives *Flavobacterium* gliding. Curr Biol 25, 338–341 (2015). 10.1016/j.cub.2014.11.045

53 Nakane, D., Sato, K., Wada, H., McBride, M. J. & Nakayama, K. Helical flow of surface protein required for bacterial gliding motility. Proc Natl Acad Sci U S A 110, 11145–11150 (2013). 10.1073/pnas.1219753110

54 Ishigami, K. et al. A Trojan horse pathogen breaking through partner-choice barriers in the insect gut. Proc Natl Acad Sci U S A 123, e2533244123 (2026). 10.1073/pnas.2533244123

55 Son, K., Guasto, J. S. & Stocker, R. Bacteria can exploit a flagellar buckling instability to change direction. Nat Phys 9, 494–498 (2013). 10.1038/nphys2676

56 Kearns, D. B. A field guide to bacterial swarming motility. Nat Rev Microbiol 8, 634–644 (2010). 10.1038/nrmicro2405

57 Abeysinghe, G. et al. Fungal mycelia and bacterial thiamine establish a mutualistic growth mechanism. Life Sci Alliance 3, e202000878 (2020). 10.26508/lsa.202000878

58 Fukuyama, S. et al. Proliferation of a bloom-forming phytoplankton via uptake of polyphosphate-accumulating bacteria under phosphate-limiting conditions. ISME Commun 5, ycaf192 (2025). 10.1093/ismeco/ycaf192

59 Schindelin, J. et al. Fiji: an open-source platform for biological-image analysis. Nat Methods 9, 676–682 (2012). 10.1038/nmeth.2019

60 Ershov, D. et al. TrackMate 7: integrating state-of-the-art segmentation algorithms into tracking pipelines. Nat Methods 19, 829–832 (2022). 10.1038/s41592-022-01507-1

